# Spatial and Temporal Inhomogeneity of Magnetic Background in Cell Culture Experiments: The Role of Type and Position of CO_2_-Incubator in a Laboratory

**DOI:** 10.1101/2023.11.14.566912

**Authors:** Ludmila M. Sencha, Maria A. Karpova, Alexey A. Dolinin, Fedor G. Sarafanov, Nikolay V. Ilin, Evgeny A. Mareev, Vladimir A. Vodeneev, Marina A. Grinberg, Irina V. Balalaeva

## Abstract

In vitro cultivation of cells in strictly controlled conditions of a biological incubator is a widely used experimental model in biological studies. The CO2-incubators allow control of temperature, atmospheric composition, and humidity; however, the recent reports point out at possible significant and incontrollable influence of incubators on magnetic background. We demonstrated that two types of CO_2_-incubators sufficiently alter the static magnetic field (SMF) in the inner chamber compared to natural geomagnetic field, and the effect depends on the incubator model. The values of SMF in the center of incubators’ chambers were lower than natural; nevertheless, the strongly expressed spatial inhomogeneity of SMF was registered, with both reduced, up to hypomagnetic, and increased SMF values. One of the incubators in operating mode generated oscillations in magnetic field with period of oscillations about several seconds and peak-to-peak amplitude depending on the measuring point within the chamber volume, up to 115% of the mean value. Since the magnetic background is considered to contribute in multiple biological effects, we emphasize the significant impact of CO_2_-incubators on magnetic background in cell culture experiments and assume that its spatial and temporal inhomogeneity may be a source for variability in cell study results.

**Highlights:**

- CO_2_-incubators sufficiently alter the static magnetic field in the inner chamber compared to natural geomagnetic field
- Spatial inhomogeneity of the magnetic field depends on CO_2_-incubator type and can reach the gradient value of more than 60 μT.
- In one type of the incubator, the generated oscillations in magnetic field were registered with a period of several seconds and a peak-to-peak amplitude up to 115% of the mean value.

## Introduction

Living organisms are constantly influenced by a plethora of chemical and physical factors. The effect of a factor on an organism can be realized through evolutionary evolved specialized molecular receptors with associated signaling pathways, or, as in the case of ionizing radiation, low-frequency electromagnetic fields, static magnetic fields, etc., predominantly through altering nonspecific molecules and supra-molecular complexes involved in various physiological processes [Binhi and Prato, 2017]. In the course of evolution, living organisms have developed adaptive mechanisms allowing their existence in the certain range of parameters of every influencing factor [Fratini and Amendola, 2014; Markov, 2005; Natan and Vortman, 2017]. Of importance, going outside the range may result in inability to respond adequately and, as a consequence, to lead to negative effects on the organism [Asashima et al., 1991; Mo et al., 2012; Satta et al., 2002].

The influence of magnetic fields (MF) on living organisms has attracted much attention of researchers in recent decades [Krylov and Osipova, 2023; Mo et al., 2014; Tian et al., 2018; Zhang et al., 2021]. The importance of the Earth’s magnetic field for normal metabolism of living organisms is well-established [Dubrov and Brown, 1978]. The geomagnetic field (GMF) varies in its magnitude and inclination angle [Finlay et al., 2010]. The GMF tends to increase from the equator (25 μT) to the north and south poles (65 μT); at the equator, the horizontal component of the field predominates with the vertical component being minimally expressed, and vice versa at the magnetic poles. In addition to geographic location, the MF in which organisms, including humans, exist is also influenced by disturbances caused by the local environment. Natural disturbances can occur due to rock deposits that have magnetic properties, and manifest as geographical magnetic anomalies [Pobachenko et al., 2016]. Also, significant disturbances to MF can be experienced by humans and other organisms in industrial and residential buildings due to elements of load-bearing wall structures and large equipment made of metals capable of magnetization. Magnetization can occur as a results of manufacturing process or long-term exposure to GMF [Gresits et al., 2015; Makinistian and Belyaev, 2018]. Another factor that disturbs the local MF are moving parts of mechanisms made of conductive materials [Mild et al., 2009].

In vitro cultivation of cells in strictly controlled conditions of a biological incubator is a widely used experimental model in biological studies. The CO_2_-incubators allow control of temperature, atmospheric composition, and humidity. However, the recent reports point out at possible significant and incontrollable impact of incubators on magnetic background due to using metal parts in the device [Gresits et al., 2015; Makinistian and Belyaev, 2018]. The altered MF can influence the cell metabolism [Sabo et al., 2002], rate of respiration [Fu et al., 2016], redox homeostasis [Zhang et al., 2017], and even proliferation and survival [Tian et al., 2018]. Moreover, the spatial inhomogeneity and temporal instability of MF in the incubator’s chamber can be the source of high variation and poor reproducibility of experimental data [Lin, 2014; Makinistian et al., 2018]. Despite the published works of several research groups, which emphasized the role of MF in cell studies, magnetic background in incubators is extremely rarely measured and can be a hidden factor that uncontrollably influences the results of cell research.

Taken into account the above, we compared magnetic field in two types of CO_2_-incubators, located in close proximity to each other. The registered MF values depended on the type and orientation of the incubator relative to GMF. The strongly expressed spatial inhomogeneity of MF was registered with both reduced and increased MF values in individual measuring points; moreover, one of the incubators generated significantly manifested temporal changes if MF value. En masse, the type and position of CO_2_-incubator in a laboratory do contribute in spatial and temporal inhomogeneity of magnetic background in cell culture experiments.

## Materials and methods

### CO_2_-incubators

MF was measured in two types of CO_2_-incubators of the same manufacturer (BINDER GmbH, Germany), located in the same laboratory room in close proximity from each other. The first type of incubators, model BINDER C150, has dimensions of 680×785×870 mm and an internal volume of 150 L. The second type of incubators, model BINDER CB53, with dimensions 400×330×400 mm has internal volume of 53 L. Both incubators have stainless steel interior lining; perforated metal shelves provided by the manufacturer also are made of stainless steel. In the basic manufacturer’s configuration, the BINDER CB53 incubator is equipped with two shelves, and the BINDER C150 is equipped with three shelves (Fig. 1A). At the bottom of the incubators’ chambers there are stainless steel trays filled with water to maintain atmospheric humidity.

**Fig. 1.**
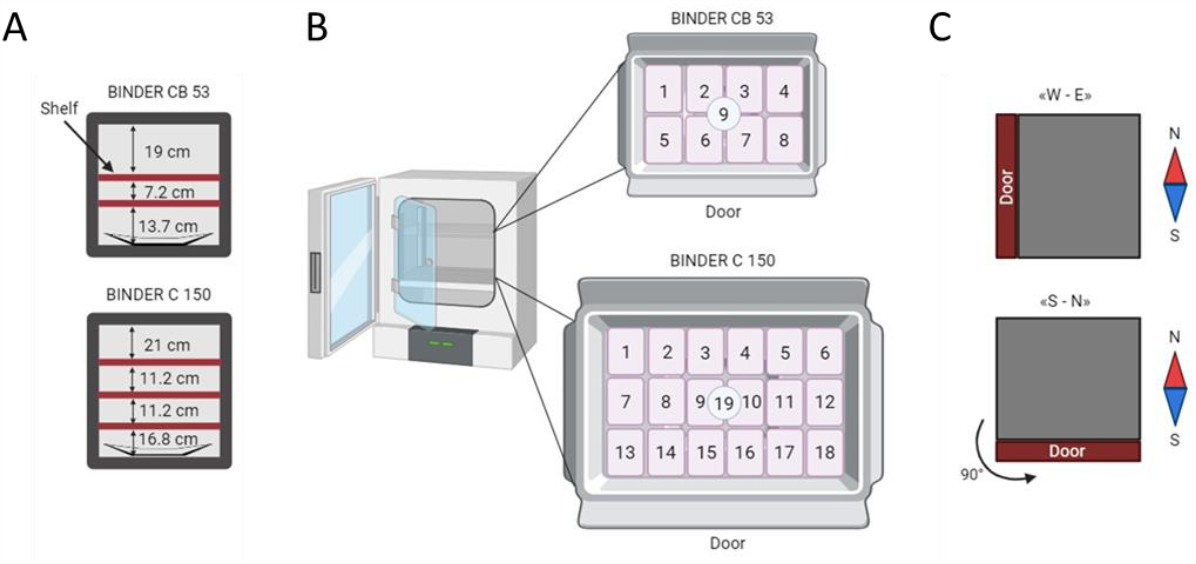
The scheme of experimental design for SMF measuring in two types of incubators, BINDER CB53 and BINDER C150. (A) Positioning of shelves in inner chambers of the incubators. (B) The location of measuring points on each shelf, corresponding to full loading of the incubators with cell culture plates. An additional measuring point was located in the center of each shelf. (C) The incubator orientation relative to the geomagnetic field when measuring in “W-E” and “S-N” positioning

### Static magnetic field measurements

Static magnetic field (SMF) measurements were performed with a Fluxgate HWT3100 3-axis magnetometer (WitMotion Shenzhen Co., Ltd., China) connected to a personal computer. Before an experiment, the sensor was calibrated according to the manufacturer’s protocol. For measuring the SMF in an incubator’s chamber, the magnetometer sensor was manually located horizontally at the measuring point on the surface of the shelf. The MF was registered for 24 s with 1 s step, followed by calculating the mean value.

The incubators were turned on 10 minutes before the start of measurements to establish a stable operating mode; the incubators’ doors during the measurements were closed. Measurements were carried out on each of the incubator shelves in positions corresponding to the locations of commonly used cell culture plates (128×85.5 mm) with the incubator fully loaded. Additionally, SMF was measured in the center of each shelf. The total amount of the measuring points was 19 in case of BINDER C150 and 9 in case of BINDER CB53 (Fig. 1B).

To assess the impact of the incubator orientation relative to the geomagnetic field, measurements were carried out at two positions of the incubator BINDER CB53: “W-E” (the incubator’s door is oriented to West) and “S-N” (the incubator’s door is oriented to South) (Fig. 1C). The incubator rotation was carried out without changing its location in the laboratory and in identical surroundings.

To assess the stability of the SMF, measurements were repeated on three different days over three weeks.

## Results

### Influence of the incubator type on the static magnetic field

The presence in the design of CO_2_ incubators of a significant number of elements with high magnetic susceptibility suggests their possible significant influence on the magnetic field (MF) in the internal chamber intended for cell cultivation. In this regard, at the first stage of the work, we compared the magnitude of the magnetic field in incubators of two types, differing in dimensions and chamber size, installed in the laboratory in close proximity to each other in the “W-E” orientation. The effect of whether the incubator was turned off or on was also assessed. For this, MF was measured for one minute before and four minutes after turning on the incubator.

The MF values measured in the center of the incubators (position 19B, middle shelf in the case of BINDER C150; position 9, bottom shelf in the case of BINDER CB53) differed significantly and in the off state were about 37 and 23 µT, respectively; the fluctuations in the field magnitude were minimal and did not exceed 0.1 µT (Fig. 2). Turning on the incubators had a very noticeable effect manifested as an appearance of MF periodical alterations. In case of BINDER C150 incubator with large chamber volume, relatively minor oscillations were recorded with a peak-to-peak value of about 2 µT (5.5% of the mean value). The period of oscillations observed by our sensor was on the order of seconds. After this incubator entered operating mode, the steady-state value of the MF was established. In the case of the BINDER CB53 incubator with a small chamber size, oscillations were significantly more pronounced, with the peak-to-peak amplitude of about 10 µT (45% of the mean value) and the period of oscillations about several seconds. Moreover, this regimen was maintained for a long observation period, more than 1 hour, which may be due to the constant operation of a rotating metal fan in the body of the device characteristic for this model.

**Fig. 2.**
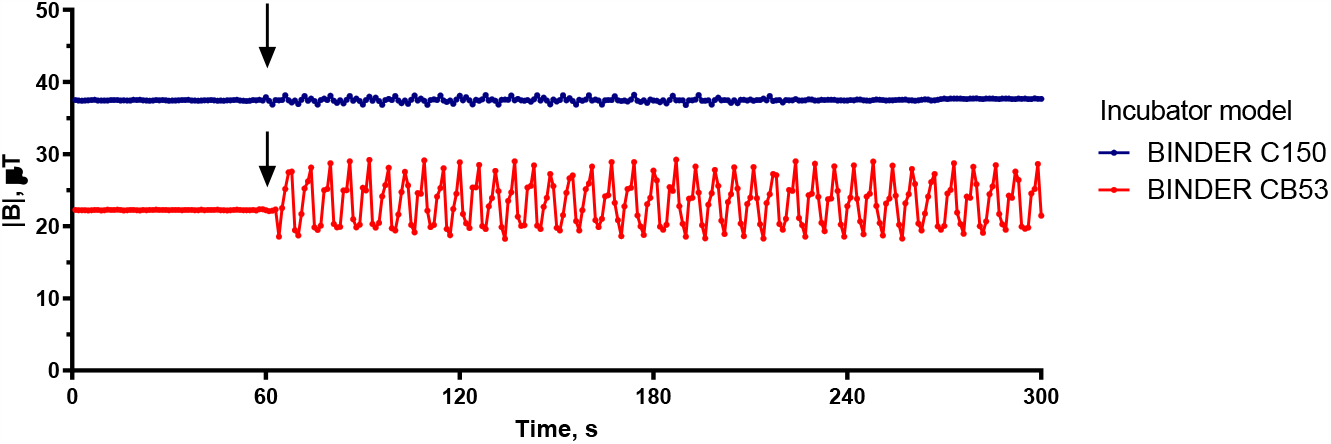
Time-dependence of MF in the center of chamber of CO_2_-incubators BINDER C150 and BINDER CB53 before and after turning on. The moment of turning the incubator on is indicated by arrow.

Based on data provided by National Oceanic and Atmospheric Administration (NOAA), Washington, USA, the modulus of the Earth’s magnetic field vector at the geographic point of the experiment (56°N, 44°E) is 53.93 µT [Chulliat et al., 2020]. The mean MF values in the centrum of both operating incubators turned out to be significantly lower than the GMF (Fig. 3). Considering the variation of the field magnitude over time, the average MF values were 37.5 ± 0.27 μT and 23.2 ± 3.18 μT in the case of BINDER C150 and BINDER CB53, respectively. It can be concluded that the magnetic field generated by the magnetized metal parts of the incubator is comparable in magnitude to GMF and has a significant effect on the ‘magnetic background’ during cell culture experiments, and the influence affects both the overall magnitude of the magnetic field and its stability over time.

**Fig. 3.**
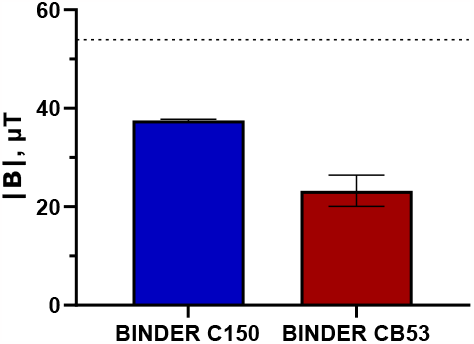
MF in the center of chamber of CO_2_-incubators BINDER C150 and BINDER CB53. The mean values were calculated as an averaged magnitude of the magnetic field within 4 min observation period; standard deviation reflects the values variability over time. The dotted line indicates the modulus of the Earth’s magnetic field at the geographic point of the experiment according to NOAA [Chulliat et al., 2020]

### Dispersion of magnetic field values over the incubator volume

Based on the assumption of the probable MF inhomogeneity inside the incubator, we measured the MF on each of the shelves in positions corresponding to the possible placement of culture plates in fully loaded incubators (Fig. 1B). The mean values during 24 s measurements were analyzed.

The data obtained indicate strong spatial variability of the magnetic field in both incubators (Fig. 4, 5). In the BINDER C150 incubator, the magnetic field values varied in the range from 9.9 μT to 72.4 μT; the peak-to-peak amplitude at a measured point was about 2-4 μT. Mapping the magnetic field in several horizontal sections of the incubator (see Fig. 4, 5A) reveals the presence of both vertical (between the lower and upper shelves) and horizontal gradients of the magnetic field. The shelf-averaged MF increased from 32.6 μT to 40.2 μT when moving from the bottom to the top shelf. On the individual shelf, the highest MF magnitude was recorded near the back wall of the chamber; when approaching the incubator door, the MF significantly decreased. The gradient “front-back” was 19.3 μT, 29.3 μT, and 37.5 μT at the bottom, middle, and top shelves, respectively. Thus, the magnitude of the gradients registered in the chamber of BINDER C150 incubator is comparable to the volume-averaged value (36.7 μT). We would like to separately note that the recorded minimum MF value of 9.9 μT can be included in the so-called hypomagnetic range (less than 10 μT) [Zhang et al., 2021]. The MF values lower than 15 μT were registered in three points of measurement; on the other hand, the region on the top shelf at the back wall of the incubator is characterized the MF values significantly exceeding the Earth’s field and reaching 72.4 μT (Fig. 4).

**Fig. 4.**
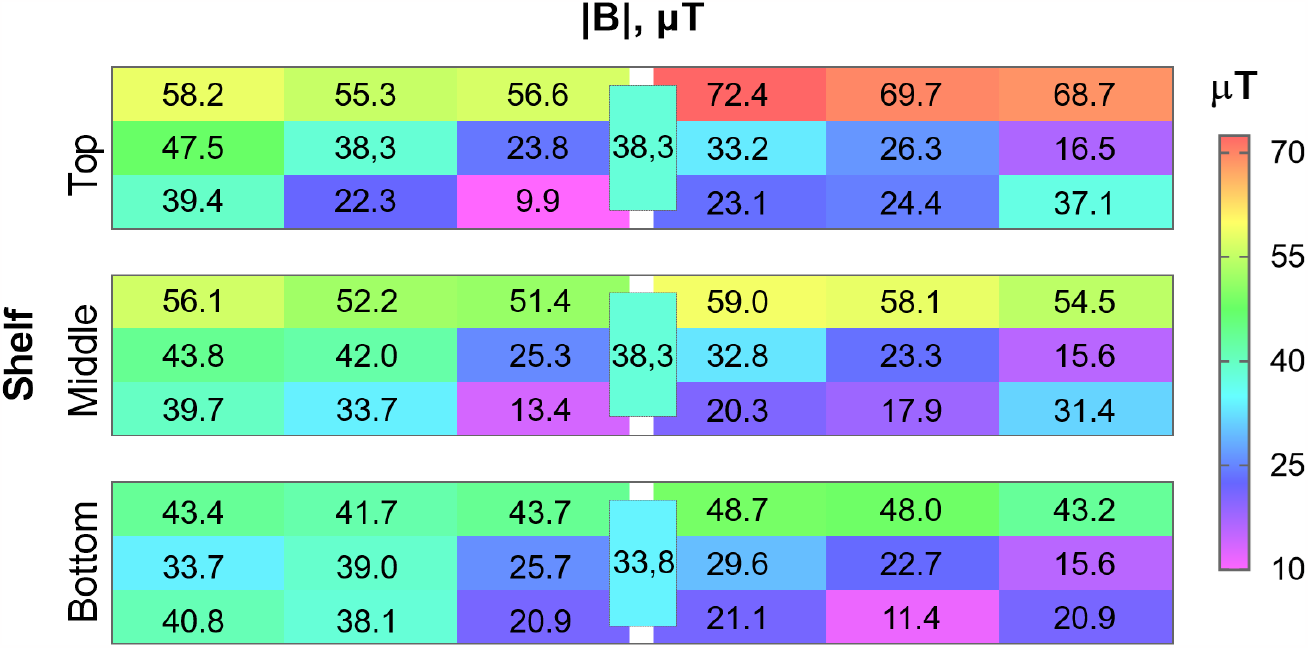
The map of MF on the bottom, middle and top shelves of BINDER C150 incubator, measured in points corresponding to location of cell culture plates at full loading of the incubator. An additional measuring point was located in the center of each shelf.

**Fig. 5.**
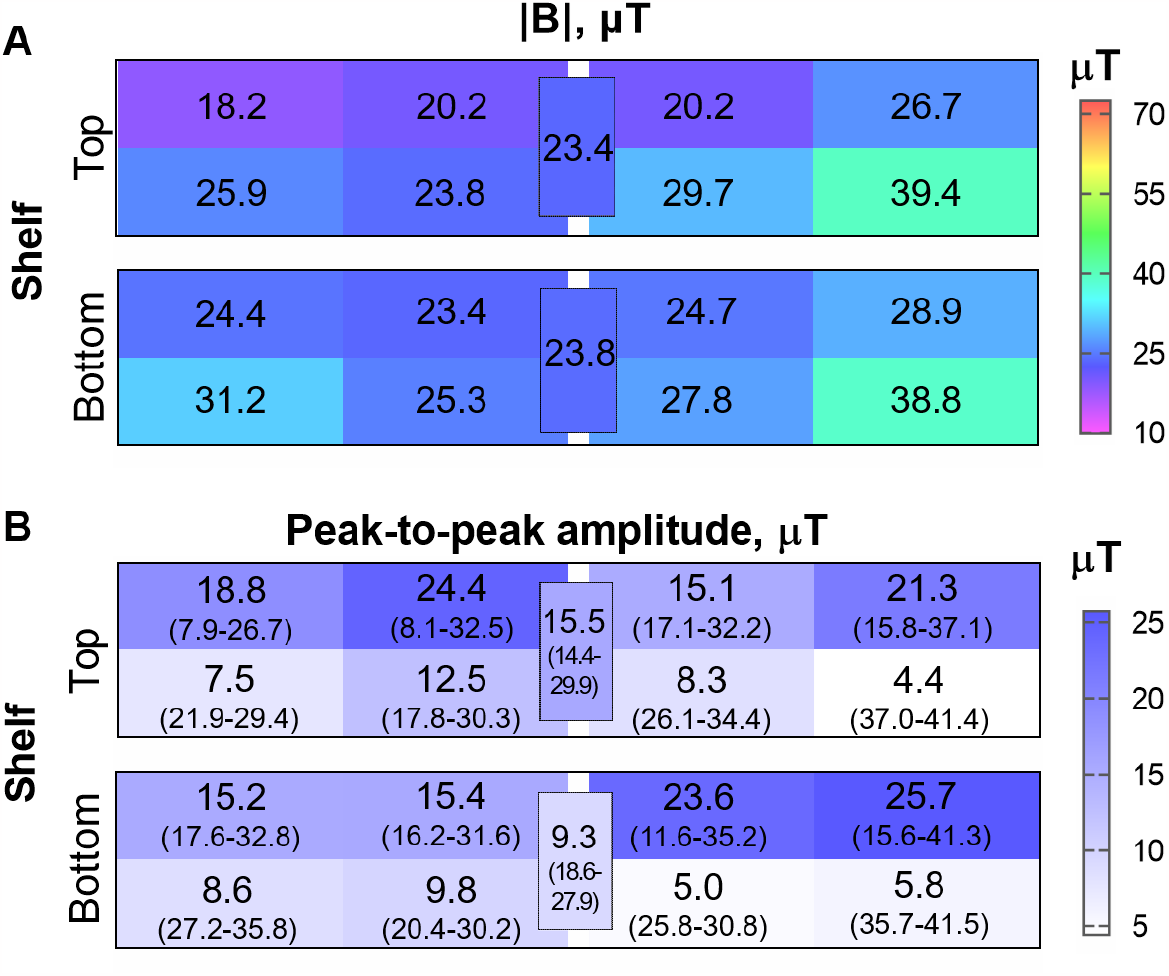
The map of MF mean magnitude (A) and peak-to-peak amplitudes (B) on the bottom and top shelves of BINDER CB53 incubator, measured in points corresponding to location of cell culture plates at full loading of the incubator. An additional measuring point was located in the center of each shelf. The range of MF variation is indicated in brackets in (B).

In case of BINDER CB53 incubator, the mean MF was relatively less variable, ranging from 18.2 µT to 39.4 µT in the chamber volume (Fig. 5A). Unlike the previous incubator, the MF near the incubator door was higher than in the depths of the chamber, with gradient “front-back” of -5.4 µT and -8.4 µT on the bottom and top shelves, respectively. The gradient between the shelves was very low, about 2.6 µT. Since we have shown the presence of significant MF oscillations for this type of incubator in operating mode, we analyzed the dispersion of the peak-to-peak amplitudes at all the measurement points (Fig. 5B). The peak-to-peak amplitudes ranged from 5 µT to 24.4 µT. The largest values were registered in the area close near the back wall of the incubator chamber, where they exceeded 50% of the mean value at almost all measuring points. Thus, the MF in this area decreased to values below 10 μT and restored to values comparable to the Earth’s magnetic field with the period of oscillations about several seconds.

In comparison, the conditions in the BINDER C150 incubator chamber are characterized by higher volume-averaged MF values and high spatial variability (36.7 ± 16 µT) compared to the BINDER C53 incubator (28.8 ± 6.6 µT) (Fig. 6). A factor that requires attention in the case of the first type on incubators is the possibility of cells being under hypomagnetic conditions or conditions of increased MF, depending on the location in the chamber. In the case of the second type of incubators, the most important are very strong fluctuations in the MF magnitude over time.

**Fig. 6.**
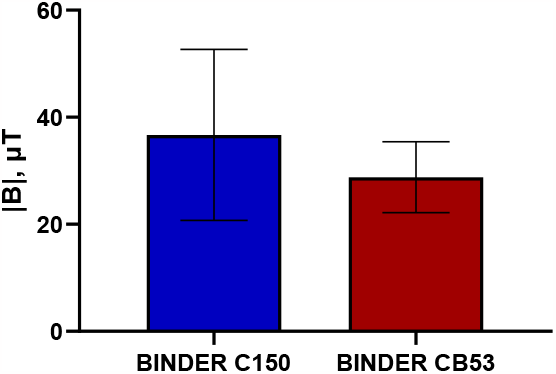
Volume-averaged MF in the chambers of CO_2_-incubators BINDER C150 and BINDER CB53. Standard deviation reflects the spatial variation of values.

To check the stability of magnetic background influencing the cells during long-term cultivation in CO_2_-incubators, we performed the repeated measurements in BINDER CB53 incubator within three-week period without changing the position of the incubator and surrounding laboratory equipment. The relative standard deviation of the mean MF magnitude did not exceed 12% based on the results of three independent measurements (Fig. 7A). However, the qualitatively different situation was observed for peak-to-peak amplitudes of MF oscillations; in several measuring points it value differed by almost two times on different days (Fig. 7B).

**Fig. 7.**
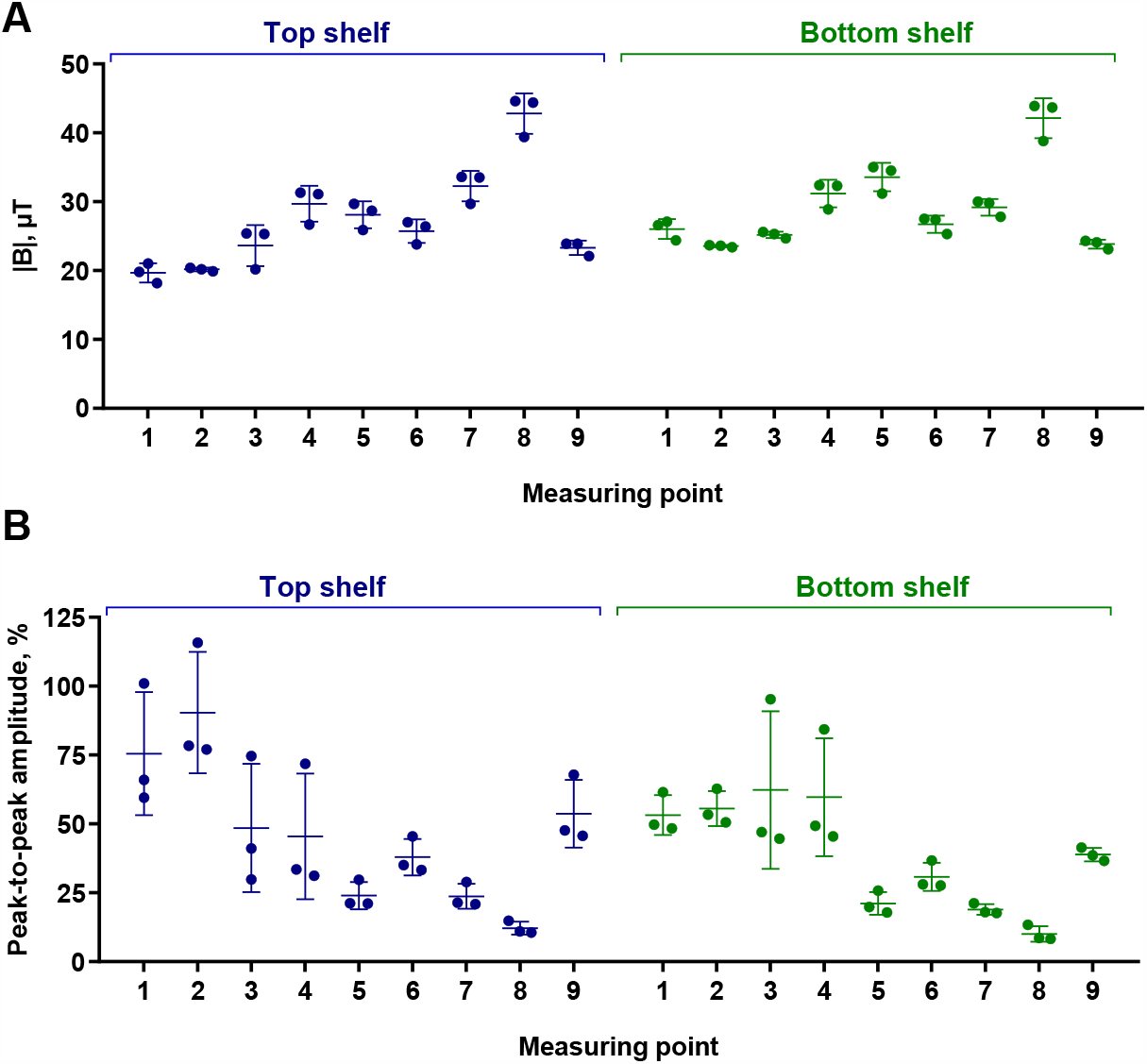
Temporal stability of the MF mean magnitude (A) and peak-to-peak amplitudes in percentage (B) in BINDER CB53 incubator over three-week period. The location of measuring points corresponds to indicated in Fig. 1. The results of three individual measurements, the mean values and standard deviation are shown.

### Influence of the incubator position in laboratory on the magnetic field

The magnitude and spatial inhomogeneity of MF in the incubator chamber are determined not only by its own structural and functional features, but also by the surroundings, including the Earth’s magnetic field, building construction elements and equipment located in close proximity. In this regard, we carried out comparative measurements of the MF in the BINDER CB53 incubator when it was rotated by 90° (changing the “W-E” to “S-N” orientation) without moving or changing the surrounding laboratory equipment.

Changing the incubator position led to significant changes in MF values in individual points; and the difference (ΔB) ranged from -16.1 µT to 9.9 µT (Fig. 8). In total, the left half of the chamber volume was characterized by maintaining the MF or increasing it, while in the right half the values significantly decreased. Such a clearly registered pattern indicates the presence of an external field source with a significant contribution to the MF inside the incubator.

**Fig. 8.**
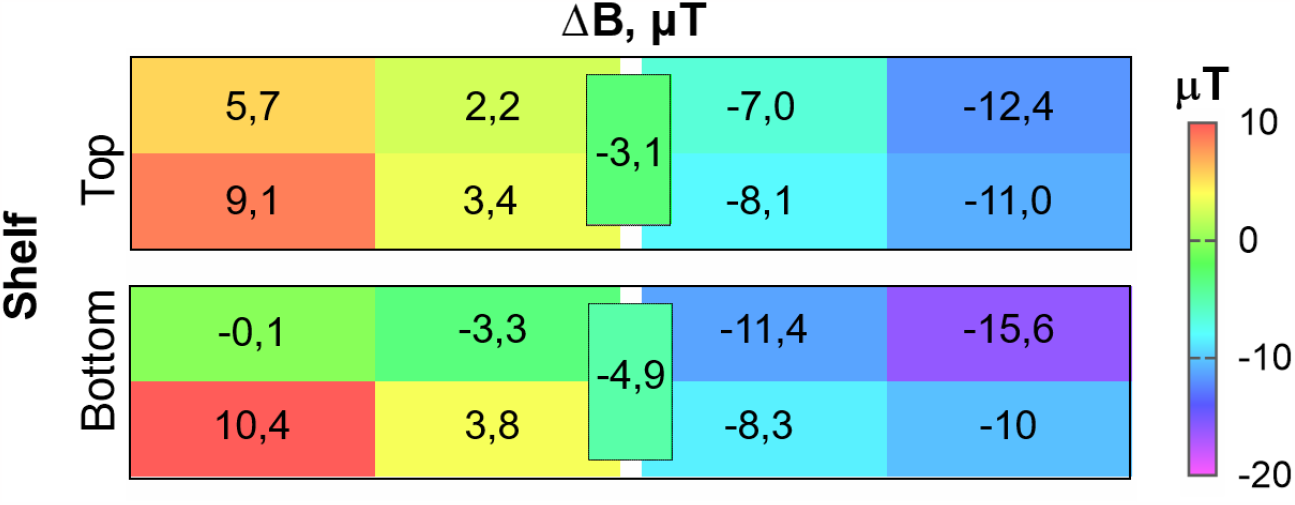
Map of changes in mean MF magnitude (ΔB) on the bottom and top shelves of BINDER CB53 incubator, induced by changing the incubator orientation from “W-E” to “S-N”. The MF was measured in points corresponding to location of cell culture plates at full loading of the incubator; an additional measuring point was located in the center of each shelf.

## Discussion

The role of magnetic field as a factor able of influencing the physiological process in living organism has been commonly accepted by the research and physician community. In recent years, a few research groups pointed out that in spite of this scientific consensus the role of magnetic background in cell culture experiments remained under-estimated whilst the altered MF magnitude in CO_2-_incubators and its inhomogeneity can be the source of significant variability and low reproducibility of the results. Thus, Portelli et al. reported the variation of MF in the chambers of different incubators in the range from < 5 µT to 450 µT, with a GMF in this region of 52.7 µT [Portelli et al., 2013]. An interesting observation was published by Mild et al., who note an increase in the MF, above the GMF values, after replacing the fan motor, despite the fact that this was an original replacement part from the manufacturer [Mild et al., 2009]. However, the number of experimental works in the field published to the moment was insufficient to attract the attention of the world scientific community, and, in our opinion, additional efforts are required to highlight the issue.

We have demonstrated that the MF in operating incubators of two types, in the center of the chamber, were significantly lower (37.5 ± 0.27 and 23.2 ± 3.18 μT) than the GMF of the region where the study was carried out (53.93 μT). However, both the incubators were characterized by high spatial MF inhomogeneity. The highest gradients were observed in BINDER C150 incubator where the MF values varied from 9.9 µT to 72.4 µT. Such magnetic field dispersion is usually not taken into account in experiments, but such heterogeneity can have an uncontrollable effect on the results. Moreover, since the cell studies usually require manipulations with multiple steps of removing the cells out of the incubator and then placing them back, changing the position of the cell plate of culture flask within the same incubator can has an additional interfering effect. The similar conclusion about high inhomogeneity has been done in a work of Makinistian and Belyaev, who demonstrated the variation from 12.3 µT to 57 µT on the shelf of the CO_2_-incubator and extremely strong impact of the holes [Makinistian and Belyaev, 2018].

The second analyzed type of incubators, BINDER C53, is characterized by lower spatial MF inhomogeneity; the MF values varied from 18.2 µT to 39.4 µT. But in this case the operation of the incubator was accompanied by the appearance of MF oscillations with the period about several seconds and magnitude comparable with the mean MF value. The oscillations were registered throughout the entire volume of the incubator chamber; in some measuring points their peak-to-peak amplitude reached more than 100% of the mean. We hypothesis that the MF oscillations registered may be caused by a rotating metal fan in the body of the device. This is a factor which has not been mentioned before as a one potentially influencing the cell study results and requiring an additional investigation.

The altered magnetic background within the CO_2_-incubators and its spatial and temporal inhomogeneity can be potentially influencing on various cellular process. It has been previously reported that hypomagnetic conditions (<10 μT) reduce cell proliferative activity [Martino et al., 2010], lower generation of reactive oxygen species [Zhang et al., 2017], modify gene transcription [Mo et al., 2014] and switch cellular metabolism to enhanced glucose consumption and glycolysis [Wang et al., 2022]. The mentioned effects were shown both on normal human or animal cells [Mo et al., 2012; Truta et al., 2012] and on tumor cells [Martino et al., 2010; Mo et al., 2016]. MF with magnitude above the GMF are also able to induce biological effects including disturbance in DNA integrity [Amara et al., 2007], induction of apoptosis [Chater et al., 2005], increased production of free radicals in cells [Vergallo et al., 2014] and reduced functional activity of macrophage cells [Flipo et al., 1998]. Despite the fact that most of the effects on activity of cells and whole organisms were studied at MF that is an order of magnitude smaller (nT) or higher (mT, T) than the MF values registered by us in CO_2_-incubators, the potential importance of magnetic background in an incubator cannot be denied. This assumption is supported, for example, by data of Martino et al. [Martino et al., 2010] who reported the statistically different proliferation rate of cells grown at 6-8 µT MF compared to 43 µT MF within the same incubator. The molecular mechanisms of how magnetic fields comparable to GMF (units to hundreds of µT) are perceived by living organisms, still needs to be elucidated.

## Conclusion

We demonstrated that two types of CO_2_-incubators sufficiently alter the static magnetic field in the inner chamber compared to natural geomagnetic field, and the effect depends on the incubator model. The values of SMF in the center of incubators’ chambers were lower than natural; nevertheless, the strongly expressed spatial inhomogeneity of SMF was registered, with both reduced, up to hypomagnetic, and increased SMF values. Changing the incubator position by its rotation leads to significant changes in SMF values distribution indicating the role of external magnetic field sources. One of the incubators in operating mode generated oscillations in MF with 5s period and peak-to-peak amplitude depending on the measuring point within the chamber volume, up to 115% of the mean value. The repeated measurements revealed the noticeable variation of mean values and especially amplitude in a three-week time period. Since the magnetic background is considered to contribute in multiple biological effects, we emphasize the significant impact of CO_2_-incubators on magnetic background in cell culture experiments and assume that its spatial and temporal inhomogeneity may be a source for variability in cell study results.

## Author contribution statement

M.A.K, L.M. S., A.A.D., F.G.S.: performed the experiments; L.M. S., M.A.K, M.A.G., A.A.D., F.G.S.: analyzed and interpreted the data; wrote the paper; I.V.B., M.A.G., V.A.V., E.A.M., N.V.I.: conceptualization and methodology; supervision and project administration, review and editing.

## References

1. Amara S, Douki T, Ravanat JL, Garrel C, Guiraud P, Favier A, Sakly M, Ben Rhouma K, Abdelmelek H. 2007. Influence of a static magnetic field (250 mT) on the antioxidant response and DNA integrity in THP1 cells. Phys Med Biol 52:4:889–898.

2. Asashima M, Shimada K, Pfeiffer CJ. 1991. Magnetic shielding induces early developmental abnormalities in the Newt, Cynops-Pyrrhogaster. Bioelectromagnetics 12:215–224.

3. Binhi VN, Prato FS. 2017. Biological effects of the hypomagnetic field: an analytical review of experiments and theories. PLoS ONE 12:e0179340.

4. Chater S, Abdelmelek H, Couton D, Joulin V, Sakly M, Ben Rhouma K. 2005. Sub-acute exposure to magnetic field induced apoptosis in thymus female rats. Pak J Med Sci 21:292–297.

5. Chulliat A, Brown W, Alken P, Beggan C, Nair M, Cox G, Woods A, Macmillan S, Meyer B, Paniccia M. 2020. The US/UK world magnetic model for 2020-2025: Technical report. DOI : 10.25923/ytk1-yx35

6. Dubrov AP, Brown FA. 1978. The geomagnetic field and life: Geomagnetobiology. New York: Plenum Press, p. 318.

7. Finlay CC, Maus S, Beggan CD, Bondar TN, Chambodut A, Chernova TA, Chulliat A, Golovkov VP, Hamilton B, Hamoudi M, Holme R, Hulot G, Kuang W, Langlais B, Lesyr V, Lowes FJ, Luhr H, Macmillan S, Mandea M, McLean S, Manoj C, Menvielle M, Michaelis I, Olsen N, Rauberg J, Rother M, Sabaka TJ, Tangborn A, Toffner-Clausen L, Thebault E, Thomson AWP, Wardinski I, Wei Z, Zvereva TI. 2010. International Geomagnetic Reference Field: The eleventh generation. Geophysical Journal International 183:1216–1230.

8. Flipo D, Fournier M, Benquet C, Roux P, Le Boulaire C, Pinsky C, LaBella FS, Krzystyniak K. 1998. Increased apoptosis, changes in intracellular Ca2+, and functional alterations in lymphocytes and macrophages after in vitro exposure to static magnetic field. J Toxicol Environ Health A 54:63–76.

9. Fratini E, Amendola R. 2014. Caves and other subsurface environments in the future exploration of Mars: the absence of natural background radiation as biology concern. Rendiconti Lincei 25:91–96.

10. Fu JP, Mo WC, Liu Y, He RQ. 2016. Decline of cell viability and mitochondrial activity in mouse skeletal muscle cell in a hypomagnetic field. Bioelectromagnetics 37:212–222.

11. Gresits I, Necz PP, Janossy G, Thuroczy G. 2015. Extremely low frequency (ELF) stray magnetic fields of laboratory equipment: a possible co-exposure conducting experiments on cell cultures. Electromagn Biol Med 34:244–250.

12. Krylov VV, Osipova EA. 2023. Molecular Biological Effects of Weak Low-Frequency Magnetic Fields: Frequency–Amplitude Efficiency Windows and Possible Mechanisms. Int J Mol Sci 24:10989.

13. Lin JC. 2014. Reassessing laboratory results of low-frequency electromagnetic field exposure of cells in culture. IEEE Antennas Propag Mag 56:227–229.

14. Makinistian L, Belyaev I. 2018. Magnetic field inhomogeneities due to CO2 incubator shelves: a source of experimental confounding and variability? R Soc Open Sci 5:172095.

15. Makinistian L, Muehsam DJ, Bersani F, Belyaev I. 2018. Some recommendations for experimental work in magnetobiology, revisited. Bioelectromagnetics 39:556–564.

16. Markov MS. 2005. «Biological windows»: a tribute to W. Ross Adey. Environmentalist 25:67–74.

17. Martino CF, Portelli L, McCabe K, Hernandez M, Barnes F. 2010. Reduction of the Earth’s magnetic field inhibits growth rates of model cancer cell lines. Bioelectromagnetics 31:649–655.

18. Mild KH, Wilén J, Mattsson MO, Simko M. 2009. Background ELF magnetic fields in incubators: a factor of importance in cell culture work. Cell Biol Int 33:755–757.

19. Mo WC, Liu Y, Bartlett PF, He RQ. 2014. Transcriptome profile of human neuroblastoma cells in the hypomagnetic field. Sci China Life Sci 57:448–461.

20. Mo WC, Liu Y, Cooper HM, He RQ. 2012. Altered development of Xenopus embryos in a hypogeomagnetic field. Bioelectromagnetics 33:238–246.

21. Mo, WC, Zhang ZJ, Wang DL, Liu Y, Bartlett PF, He RQ. 2016. Shielding of the geomagnetic field alters actin assembly and inhibits cell motility in human neuroblastoma cells. Sci Rep 6:22624.

22. Natan E, Vortman Y. 2017. The symbiotic magnetic-sensing hypothesis: do Magnetotactic Bacteria underlie the magnetic sensing capability of animals? Mov Ecol 5:1–5.

23. Pobachenko SV., Shitov AV, Grigorjev PE, Sokolov MV, Zubrilkin AI, Vypiraylo DN, Solovjev AV. 2016. EEG reactions of the human brain in the gradient magnetic field zone of the active geological fault (pilot study). Izvestiya, Atmospheric and Oceanic Physics 52:745–752.

24. Portelli LA, Schomay TE, Barnes FS. 2013. Inhomogeneous background magnetic field in biological incubators is a potential confounder for experimental variability and reproducibility. Bioelectromagnetics 34:337–348.

25. Sabo J, Mirossay L, Horovcak L, Sarissky M, Mirossay A, Mojzis J. 2002. Effects of static magnetic field on human leukemic cell line HL-60. Bioelectrochemistry 56:227–231

26. Satta L, Antonelli F, Belli M, Sapora O, Simone G, Sorrentino E, Tabocchini MA, Amicarelli F, Ara C, Cerù MP, Colafarina S, Conti Devirgiliis L, De Marco A, Balata M, Falgiani A, Nisi S. 2002. Influence of a low background radiation environment on biochemical and biological responses in V79 cells. Radiat Environ Biophys 41:217–224.

27. Tian X, Wang D, Zha M, Yang X, Ji X, Zhang L, Zhang X. 2018. Magnetic field direction differentially impacts the growth of different cell types. Electromagn Biol Med 37:114–125.

28. Truta Z, Truta M, Micu R. 2012. Zero magnetic field influence on human spermatozoa glucose consumption. Rom Biotechnol Lett 17:6928–6933.

29. Vergallo C, Ahmadi M, Mobasheri H, Dini L. 2014. Impact of inhomogeneous static magnetic field (31.7–232.0 mT) exposure on human neuroblastoma SH-SY5Y cells during cisplatin administration. PLoS ONE, 9:e113530.

30. Wang GM, Fu JP, Mo WC, Zhang HT, Liu Y, He RQ. 2022. Shielded geomagnetic field accelerates glucose consumption in human neuroblastoma cells by promoting anaerobic glycolysis. Biochem Biophys Res Commun 601:101–108.

31. Zhang Z, Xue Y, Yang J, Shang P, Yuan X. 2021. Biological effects of hypomagnetic field: Ground-based data for space exploration. Bioelectromagnetics 42:516–531.

32. Zhang HT, Zhang, ZJ, Mo WC, Hu PD, Ding HM, Liu Y, Hua Q, He RQ. 2017. Shielding of the geomagnetic field reduces hydrogen peroxide production in human neuroblastoma cell and inhibits the activity of CuZn superoxide dismutase. Protein Cell 8:527–537.

